# Correlations across timing cues in natural vocalizations predict biases in judging synthetic sound burst durations

**DOI:** 10.1101/2022.05.29.493898

**Authors:** Matthew Jané, Sashank Pisupati, Kasey E. Smith, Luan Castro-Tonelli, Liana Melo-Thomas, Rainer K.W. Schwarting, Markus Wohr, James J. Chrobak, Heather L. Read

**Affiliations:** Department of Psychological Sciences - Behavioral Neuroscience Division, University of Connecticut; Princeton Neuroscience Institute, Princeton University; Behavioral Neuroscience, Experimental and Biological Psychology, Faculty of Psychology, Phillips-University of Marburg; Center for Mind, Brain and Behavior, Phillips-University of Marburg; Faculty of Psychology and Educational Sciences, Research Unit Brain and Cognition, Laboratory of Biological Psychology, Social and Affective Neuroscience Research Group, KU Leuven; Leuven Brain Institute, KU Leuven; Department of Biomedical Engineering, University of Connecticut

## Abstract

It is well known that animals rely on multiple sources of information in order to successfully identify sounds in natural environments, to make decisions that are optimal for their survival. For example, rats use duration and pitch cues to respond appropriately to prosocial and distress vocalizations (Saito et al., 2019). Vocalization duration cues are known to co-vary with other temporal cues (Khatami et al., 2018), yet little is known about whether animals rely upon such co-variations to successfully discriminate sounds. In the current study, we find natural alarm vocalizations in rats have onset and offset slopes that are correlated with their duration. Accordingly, vocalizations with faster onset slopes are more likely to have shorter durations. Given that vocalization slopes begin and end within milliseconds, they could provide rapid perceptual cues for predicting and discriminating vocalization duration. To examine this possibility, we train rodents to discriminate duration differences in sequences of synthetic vocalizations and examine how artificially changing the slope impacts duration judgments. We find animals are biased to misjudge a range of synthetic vocalizations as being shorter in duration when the onset and offset slopes are artificially fast. Moreover, this bias is reduced when rats are exposed to multiple synthetic vocalization bursts. The observed perceptual bias is accurately captured by a Bayesian decision-theoretic model that utilizes the empirical joint distribution of duration and onset slopes in natural vocalizations as a prior during duration judgements of synthetic vocalizations. This model also explains why the bias is reduced when more evidence is accumulated across multiple bursts, reducing the prior’s influence. These results support the theory that animals are sensitive to fine-grained statistical co-variations in auditory timing cues and integrate this information optimally with incoming sensory evidence to guide their decisions.

## Introduction

When someone jams their toe on a door and belts out an expression of, “oww, oww, oww”, how do you judge whether they are truly hurt or just mildly annoyed? Certainly, the duration, loudness, and pitch of each “oww” will help you judge the expression (Belin et al., 2008; Jürgens et al., 2018; Lausen and Hammerschmidt, 2020). The slope of sound level increase with the onset and offset of each “oww” is also an important cue (Cumming et al., 2015; Paquette and Peretz, 1997; Stecker and Hafter, 2000; Grassi and Darwin, 2006; Friedrich and Heil, 2017). To make an accurate judgment, it could be important to hear “oww” repeated multiple times to accumulate sound feature information (Brunton et al., 2013). Finally, there can be unique combinations of temporal features in sounds that convey the information needed to properly judge and categorize them (Bizley and Cohen, 2013). Accordingly, the effective judgment of such expressions could require detecting a combination of acoustic features and their co-variations on multiple timescales.

Though much is known about how acoustic features themselves shape perception, far less is known about how their statistical variations may do so. Previously, we observed a physical limit in vocalization sequences that humans and other animals generate where longer vocalizations cannot be repeated faster than the duration allows (Khatami et al., 2018). This results in an increase in the statistical probability that long-duration vocalizations will have slower repetition rates than short-duration vocalizations. In other words, the repetition rate could be predictive of vocalization duration or vice versa. Indeed, mathematically, the combination of vocalization duration and repetition rate objectively differentiates vocalization type or category across a wide range of animals including humans (Khatami et al., 2018). On shorter timescales, humans rely heavily on the rate or slope of sound onset to judge the duration or loudness of sound (Stecker and Hafter, 2000; Grassi and Darwin, 2006; Friedrich and Heil, 2017). Indeed, it has been suggested that the sound onset slope is a more informative cue for sound duration than the sustained sound duration itself (Friedrich and Heil, 2017)! However, there are no theories for how such cue interactions come about. One well-supported theory is that perception is strongly influenced by statistical variations of acoustical cues found in natural sounds (Elliott and Theunissen, 2009; Geffen et al., 2011; McDermott and Simoncelli, 2011; Zhai et al., 2020) Along these lines, we propose that statistical co-variations between onset and duration in natural sounds could explain why sound onsets strongly influence the perception of sound duration.

Here we explore and computationally model how animals judge sound durations with independent variation in sound slope. We examine these perceptual interactions in rodents, as they share similar brain organization and sound duration perception with other mammals including humans (Kelly et al., 2006; Read and Reyes, 2018). Here, rodents are trained to judge duration differences in large sets of synthetic sound burst sequences with durations ranging from 100 to 250 ms. The durations of these sound burst sequences are similar to those found in natural rodent vocalizations. However, our sound design allows us to artificially impose fast slopes on longer-duration sounds to explore how sound slope impacts the perception of duration. Intriguingly, we find long-duration sounds with faster than normal slopes are systematically misjudged as being shorter in duration. This perceptual misjudgment or bias dominates when only a single sound burst is heard; whereas, a more accurate judgment prevails when multiple sound bursts are heard. We find that the observed misjudgments are well explained by a Bayesian model of decision-making that incorporates “prior experience” with natural vocalization statistics into synthetic vocalization judgements. Specifically, since the onset and duration are negatively correlated in natural vocalizations, incorporating this prior into duration judgements about synthetic vocalizations introduces a bias towards shorter durations when slopes are artificially fast. This model also accurately captures the improved performance and decreased bias seen in the behavior when more sensory evidence is accumulated over repeated sound bursts. These results support the idea that sound duration judgments reflect optimal integration of prior experience with ongoing accumulation of sensory information.

## Materials and Methods

### Quantifying Statistical Variations of Temporal Cues in Natural Alarm Vocalizations

Prior studies found that rats and humans readily discriminated sound durations greater than 100 milliseconds long (Kelly et al., 2006) and that the slope of sound onset altered duration perception (Cumming et al., 2015; Paquette and Peretz, 1997; Stecker and Hafter, 2000; Grassi and Darwin, 2006; Friedrich and Heil, 2017). However, the statistical variations and relationships between slope and duration temporal cues have not been described for natural sounds. Here, we examined the statistical distributions of onset, offset, and duration temporal cues found in natural rodent alarm vocalizations (Fig. 1). Alarm vocalizations were generated by rodents in a conditioning paradigm as described previously (Melo-Thomas et al., 2020). and the vocalizations made in the absence of haloperidol were used and are available in an online data repository (DOI: 10.5281/zenodo.5762778). Here, 1330 vocalizations were selected for analysis based on having a spectral center of mass around 22kHz (24.7, ± 0.87), as is characteristic of alarm vocalizations (Fig. 1A). Additionally, two raters screened each vocalization to make sure all artifacts were removed from the analysis. Both raters, who were blind to the decisions of the other rater, showed high levels of agreement (intra-rater percent agreement = 99.4%, inter-rater percent agreement = 98.0%). The onset, offset and intervening duration cues of each vocalization were determined using an approach detailed previously (Khatami et al., 2018). Briefly, a Hilbert transform was performed to recover the positive sound envelope (Fig. 1B, Teal line). Vocalization onset was defined as that point where the envelope sound level rose to 10 standard deviations above the noise floor (Fig. 1C,*t*_onset_). Vocalization offset was determined as a return to the noise floor or baseline sound level (Fig. 1C, *t*_offset_). The amplitude of each call was set as a ratio of the area under the curve (*AUC*) of the envelope (*Y* (*t*)) of the respective call such that,

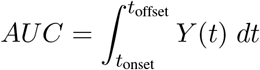

**Figure 1:**
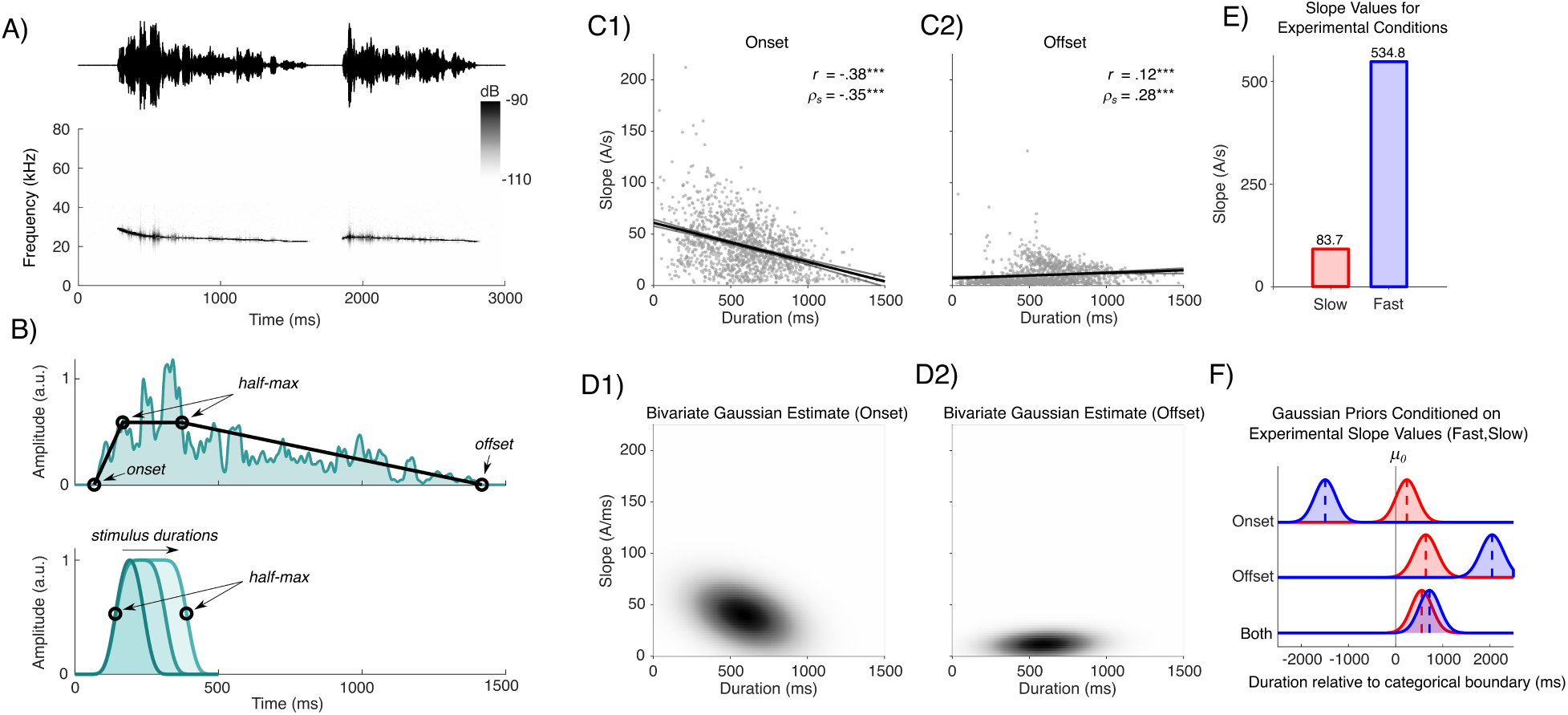
Analysis of natural vocalizations and correlation of sound onset/offset slope and plateau duration. A) Example of raw 22kHz vocalizations collected from one rat recording with the corresponding spectrogram. The examples shown have a center of mass at 25.0 kHz. B) [upper panel] Envelope of the first vocalization in panel A. Points show the time of sound onset, time of offset, and the time at the half-max values. [lower panel] Synthetic vocalizations used in discrimination task for the slow condition (short, middle, longest duration conditions). C1 and C2) Onset and offset slopes systematically co-vary with vocalization duration. Scatter plots illustrate the vocalization duration and slopes for each of the 1330 vocalizations analyzed (Methods). Vocalization durations ranged from 28 to 1467ms. C1) Onset slope decreases as vocalization duration increases following resulting in a strong negative (inverse) correlation. Pearson product-moment (r) and spearman rank-order (*ρ*_*s*_) correlations are displayed with a significance level (*** p*<*.0001). C2) Offset slope increases as vocalization duration increases resulting in a weak positive correlation. D) [panel 1] Estimated bivariate Gaussian for duration and onset slope based on data from panel C1. [panel 2] Estimated bivariate Gaussian for duration and offset slope based on data from panel C2. E) Showing the slope values of the experimental slope conditions based on the stimulus duration at the categorical boundary (175 ms). F). Conditional univariate Gaussians derived from multivariate Gaussians. Conditional values come from the slope values in panel E. The distributions corresponding to “joint” are derived via a trivariate gaussian where duration, onset slope, and offset slope comprise the three axes.

In order to calculate onset and offset slope, we first calculated the change in amplitude and the change in time for each vocalization in the data set. The change in relative amplitude value was mathematically be defined as:

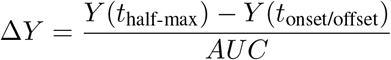

Vocalization “plateau” duration was quantified as the time between the vocalization half-maximum following onset and the vocalization half-maximum prior to offset (Fig. 1C). In our set of 1330 vocalizations, plateau duration varied from 28 to 1467ms with a median and mean duration of 581 and 586ms. The vocalization onset slope was quantified by calculating the absolute amplitude rate of change between vocalization onset (*t*_onset_) and the first half-maximum peak (*Y* (*t*_half-max_)). Likewise, the vocalization offset slope was quantified as the amplitude change between vocalization offset (*t*_offset_) and the first half-maximum peak (*t*_half-max_). Finally, the absolute values were used to quantify the average onset and offset slope for each vocalization as summarized (Fig. 1E).

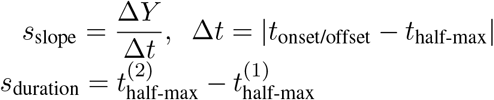

### Synthetic Vocalizations for Perceptual Testing

To behaviorally test the effects of sound slope on duration perception, we generated synthetic vocalizations with a subset of slope and duration temporal cues observed in natural vocalizations (Fig. 1). In behavioral tests (Fig. 2), animals judged a set of seven synthetic vocalizations durations as being short or long in duration (Fig. 2C). The seven synthetic vocalizations ranged in duration from 100 to 250 milliseconds (Fig. 2C) and fell within the range of alarm vocalization plateau durations reported above and summarized as a joint scatter plot (Fig. 1 C1). For synthetic vocalizations, the onset and offset slopes were the same (symmetric) for all 7 sound durations used in behavioral testing. Synthetic vocalization durations were defined by a square wave sound pressure waveform envelope. Accordingly, square wave sound pulses defined 7 different plateau durations (100, 130, 160, 175, 190, 220, and 250 ms) spanning the lower end of the natural vocalization duration range of 100 to 250 ms.

**Figure 2:**
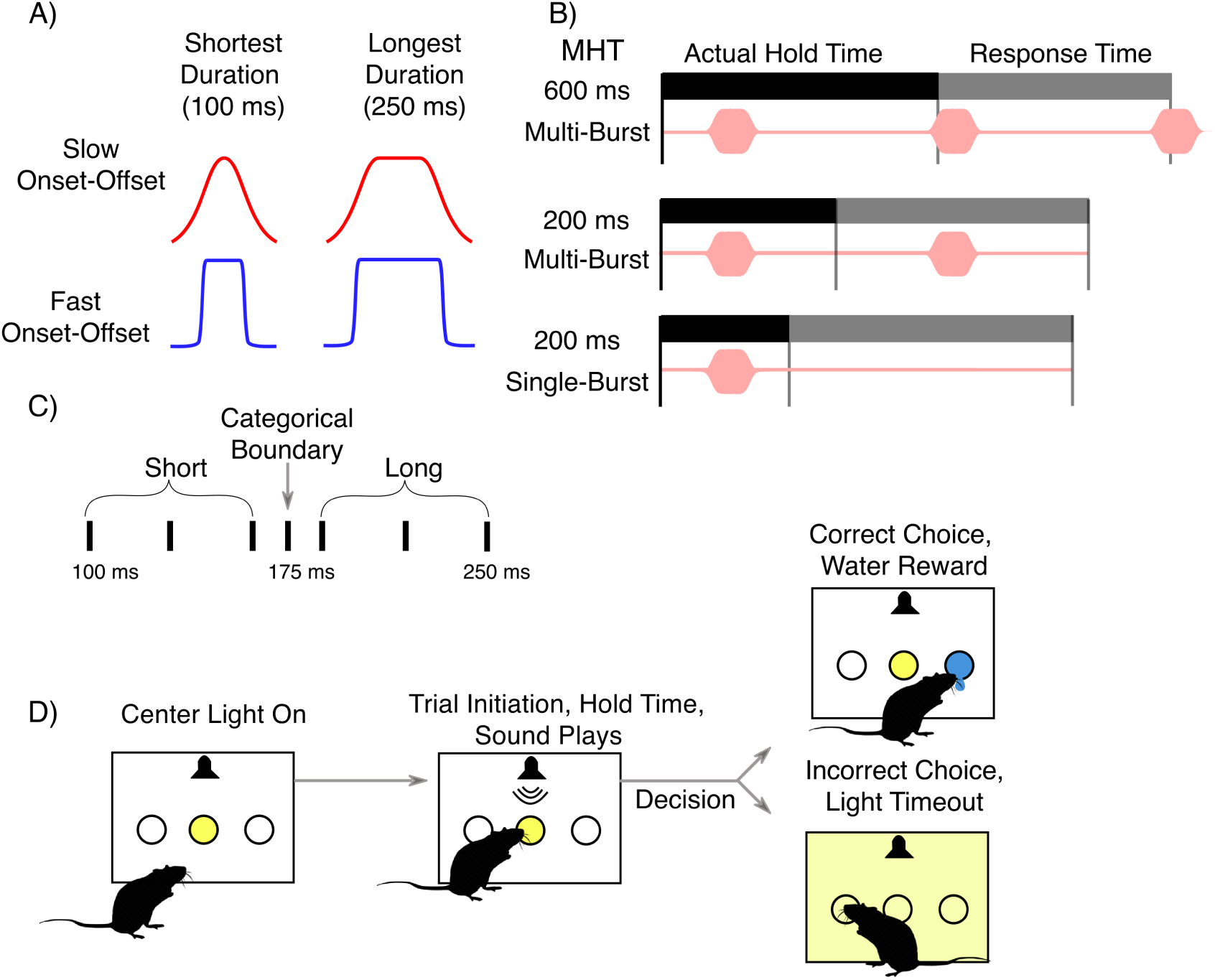
Experimental design. A) An illustration of sound envelopes for the two sound slope conditions (red = slow; blue = fast) and the shortest and longest plateau sound durations. In all variations of the behavioral tasks, animals judged short versus long duration for seven different sound bursts varying in plateau durations between 100 and 250 ms. B) Three example single trials to illustrate the relationship between sound burst sequence played (pink bursts), the required minimum hold time (MHT) condition (top, middle, bottom row), and the actual hold time at the center port (black verticle line). Across all trials and animals, the average proportion of bursts heard during the actual hold time increases with the hold time and MHT as shown in Figure 3C-D. C) Experimental layout of incremental sound plateau duration showing the duration of sounds that are played within a given session. Rats are rewarded for choosing left if the stimulus duration is greater than 175ms and rewarded for choosing right if the stimulus duration is less than 175ms. D) Sequence of events in a given trial. The animal pokes the center port to initiate the trial and the playing of sounds, then they must hold their nose in the center port for the minimum hold time depending on the MHT condition (see panel B). Then the rat makes a decision by poking their nose into the right or left port. In the case of a correct choice the rat receives 25mL of water, while in the case of an incorrect choice the rat receives a 30 second time out light which does not let them start a new trial for 30 seconds.

The fast and slow slopes were chosen to span the extreme ends of natural vocalization distribution (Fig. 2A, red and blue lines, respectively). To vary the slope of synthetic vocalizations, the square wave sound pulses were smoothed with a Basis spline (B-spline) filter function, as detailed previously (Lee et al., 2016). In two separate sets of sounds, the B-spline cutoff frequency was either 5 or 32 Hz to generate slow (83.7 A/s) versus fast (534.8 A/s) onset-offset slopes, respectively. The average slow onset-offset slope (Fig. 1E, red bar, 83.7 A/m) used for behavioral testing fell within the range of onset slopes found in natural alarm vocalizations (Fig. 1C1). In contrast, the fast onset-offset slopes were more than 2 fold faster than the fastest onset slopes observed for comparable duration alarm vocalizations (Fig. 1E, blue bar; Fig. 1C1). This allowed for high cue contrast with the average fast onset-offset slope (Fig. 1E, blue bar) being 6.4 fold faster than the average slow onset-offset slope (Fig. 1E, red bar). The corresponding average fast and slow slopes were 534.8 A/ms versus 83.7 A/ms, respectively (Fig. 1E). Synthetic onset and offset sound slopes were estimated as the absolute approximate derivative at the first and second half-max points of the sound envelope, respectively. Given that onset and offset slopes were symmetric, *s*_slope_ both slopes were defined by the following equation.

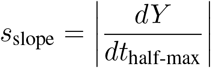

Accordingly, square wave sound pulses defined 7 different plateau durations (100, 130, 160, 175, 190, 220, and 250 ms) spanning the lower end of the natural vocalization duration range of 100 to 250 ms. These square wave sound pulses were smoothed with a Basis spline (B-spline) filter function having a cutoff frequency of either 5 or 32 Hz to generate slow (83.7 A/s) and fast (534.8 A/s) onset-offset slopes, respectively. For each burst duration, the total energy of the sound was adjusted to be equal across the two onset-offset sound conditions, to minimize the saliency of this cue. The onset times for any single sound burst in a sequence were staggered over a 125 ms window to approximate the average 2 Hz sound burst rate found in the 22 kHz vocalization sequences. The latter approach allowed us to minimize perceptual reliance on periodicity cues. For the 7 different sound durations, there were 100 different sequence variations and a total of 700 sound sequences for each onset-offset slope type. Thus, there were 1400 different sound burst sequences for the two onset-offset slope conditions. For all sequence variations, each sound burst had a unique random combination of tonal frequencies to reduce reliance on pitch perception for sound judgment.

Our synthetic vocalizations differed from natural vocalizations in several key ways. Our synthetic vocalizations were devoid of pitch cues, so we could probe temporal cue sensitivities. For all sequence variations, each sound burst had a unique random combination of tonal frequencies to reduce reliance on pitch perception for sound judgment. Thus, instead of having a fundamental frequency of 22 kHz with a harmonic frequency at 44 kHz, the synthetic vocalizations were shaped white noise. Natural vocalizations had different (asymmetric) onset versus offset slopes (Fig. 1B upper plot) but our synthetic vocalizations had the same (symmetric) onset and offset slope (Fig. 1B, lower plot). With this symmetry, shorter inter-vocalization intervals were possible allowing us to test sensitivity to slope and duration over more trials in a given block. In natural vocalizations, the onset and offset slopes were negatively or positively correlated with vocalization duration, respectively (Fig. 1C1 and C2, respectively). Moreover, onset slope and duration were more strongly correlated than the offset slope and duration (Fig. 1C1). Accordingly, the onset slope versus duration correlation coefficient was -.38 (p*<*0.0001, N=1330 vocalizations) and the offset slope versus duration correlation coefficient was 0.12 (p*<*0.0001, N=1330). In our synthetic vocalizations, the slope and duration did not co-vary (correlate) across the 7 different sound durations. Instead, for a given set of sounds (e.g. the fast slope sounds) the slopes were the same across all 7 sound durations. This allowed us to determine if the artificially imposed fast or slow sound slopes would uniformly shift the perception of duration.

### Using Natural Vocalization Statistics to Define the Prior of a Bayesian Model of Duration Judgement

Though prior studies examined how sound onset slope impacts loudness and duration perception, no theory for how such cue interactions come about had been formulated. Here, we hypothesized that onset slope impacts sound duration perception because the two temporal cues co-vary in natural sounds such as alarm vocalizations. To address this hypothesis, we developed computational models based on natural sound statistics to predict shifts in duration judgment behavior observed with changes in slope. As detailed above, we quantified the probabilities of three cue distributions found in natural alarm vocalizations including onset, offset, and duration. Next, we quantified the co-variations or correlations in these cues including 1) the joint distribution of onset slopes and durations (”onset” prior type), 2) joint distribution of offset slopes and durations (”offset” prior type), and 3) the three-way joint distribution of onset slopes, offset slopes, and durations (”both” prior type). Finally, we used the Gaussian approximations of the three joint probability distributions as conditional priors in our Bayesian models to simulate the behavioral judgment of synthetic vocalizations under our three task conditions.

Our first two steps to building our Bayesian model included quantifying the probabilities and correlations between slope and duration temporal cues found in natural vocalizations. To quantify the joint distribution of onset-offset slopes and duration cues found in natural alarm vocalizations, we fit a 2-D Gaussian to the empirical joint distributions of slopes and duration for 22 kHz alarm calls. The *μ* (2d vector of means) and Σ (2×2 covariance matrix) are the maximum likelihood estimates of slope and duration parameters in the joint probability distribution. The mean probabilities of the fast and slow slope conditions were then used as Gaussian priors in our Bayesian models (Fig. 1F). To incorporate these natural sound statistics into our Bayesian model, the maximum likelihood estimation was utilized to fit a multivariate Gaussian (bivariate for onset and offset prior types, trivariate for both prior type) as follows:

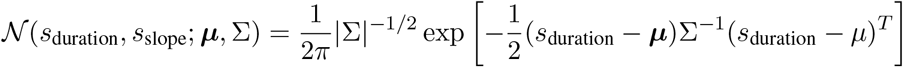

To calculate conditional prior distributions, the slope (*s*_slope_) is set to the experimental slope conditions for slow (83.7 A/ms) and fast (534.8 A/ms).

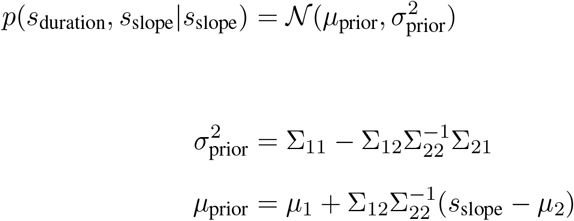

These two Gaussians served as priors in our Bayesian Decision theoretic model used to simulate and in a sense predict the sound duration judgement behavior, as detailed below.

### Automated Behavioral Training and Testing System

Rats performed all behavioral tasks inside an acrylic crate located within a single-walled sound isolation chamber. Three nose ports containing photodiode sensors were located on the back wall of the acrylic box. An ultrasonic speaker (Avisoft Bioacoustics) was located along the back wall of the sound isolation chamber at approximately 7 cm above and 18 cm in front of the center nose port. Based on nose-poke behavior and computer-generated task conditionals, water reward was delivered automatically at a rate of 6 mL/min through Teflon tubing (17 gauge) located at the center-left and right nose ports. Reward volume varied with the phase of training but was approximately 25-50 μL for each correct choice. Behavior was monitored, sound, light, and water delivery was controlled by custom MATLAB software (Mathworks, Natick MA), in conjunction with Arduino-based pulse generator and state machines (Sanworks).

### Flexible Perceptual Categorization Task: Initial Training

To determine how onset-offset slope and task conditions impact perception of sound duration, we trained male Long Evans rats (from Envigo) to perform a flexible perceptual categorization in a binomial choice task (Jaramillo and Zador, 2014). All animal procedures were approved by the Institutional Animal Care and Use Committee (IACUC) at the University of Connecticut.

First, animals were acclimated to a reverse day-night cycle for training and testing and learned to obtain daily water by poking their nose into the nose ports to receive water reward. Animal weights were monitored so that they did not fall below 80% of the individual’s baseline.

To learn the binomial choice task, animals were progressed through six training phases. In phase 1, animals were acclimated to hearing sound stimuli and obtaining their daily water allotment by poking their nose in any one of the three nose ports to release a water reward. In phase 2, sequences of the shortest (100 ms) or longest (250 ms) duration sound bursts were played each time animals held their nose in the center nose port for 150 ms. The sound sequence for that trial would continue to play until the animal by chance poked their nose in the appropriate left or right side-port associated with long and short duration sounds, respectively. In Phase 3, the required minimum hold time (MHT) for holding and hearing sound at the center nose port was increased from 150 ms to 600 ms in 2 ms increments per trial. A bright overhead light delivered a cue for a 6-second timeout when animals failed to hold for the MHT. During this timeout, rats were unable to initiate a new trial in the center port. Phase 4, additionally required that rats respond (choose a side port) within 4 seconds after the MHT. In phase 5, the overhead light was a cue for a 30-second timeout when animals choose the incorrect side port for the 100 and 250 ms duration sounds. This ended the trial and required rats to start a new trial b. Phase 5 was completed when animals correctly judged long (250 ms) and short (100 ms) duration sounds with an average percent correct of 77%. In Phase 6, animals learned to judge 100 and 250 ms sound durations as well as 5 additional intermediate sound durations. Generally, throughout all phases, rats were moved to each new phase following sessions of ≥ 115 correct hold trials.

Since animals can develop motor biases toward choosing the left or right side, several anti-biasing measures were employed throughout all phases of training. First, no more than 3 trials of the same plateau duration were presented in a row. Additionally, after every 25 trials, a custom-written Matlab program automatically evaluated the animals’ bias and increased trial numbers as well as the reward volume (by 50 uL) or correct choices on the side opposite of the bias.

### Final Training & Testing

In Phase 6, our goal was to compare the ability to judge sound durations with either fast or slow onset-offset cues under similar conditions. During phase 6, animals judged sound durations with fast versus slow onset-offset slopes in alternating blocks or “test sessions” with a fixed onset-offset slope condition. For inclusion in our final analysis, a single session needed to have ≥115 correct hold trials, ≥ 77% correct choice performance on the cardinal duration sound bursts, and show no significant bias. Rats completed 8 sessions meeting the criteria (4 at each onset-offset slope condition) with a required hold time of 600 ms.

To test the impact of reducing the accumulation of sound bursts heard, we introduced two additional conditions. First, we reduced the MHT from 600 to 200 milliseconds and animals continued to hear multiple sound bursts until they made a choice. Next, the hold time was maintained at 200 ms and only a single burst sound burst was played for animals to judge duration. On average, 21 and 23 training sessions (days) were required for animals to meet the criteria for the 600 and 200 hold time conditions, respectively. On average, 37 sessions (days) were required for animals to reach the training criterion to judge the single burst sound condition.

### Estimated sound bursts accumulated based on hold time

Since the sound sequences continue to play throughout the MHT until animals make a choice, the total number of sound bursts heard during the hold time was reduced by reducing the MHT from 600 to 200 ms and from multiple bursts to a single burst (Fig. 3). In our binomial choice task, animals were free to remain at the center port for longer before making their side port choice and sound continues to play until they made a choice. Thus, animals accumulated sound evidence during the “hold time” between initiating and completing a trial with a final choice at a side port. The actual hold times were all longer than the MHT and varied with the multi-burst versus single burst conditions (Fig. 3D) but minimally with the sound slope conditions (Fig. 3D, red vs blue symbols). We estimated the proportion of sound bursts heard for each task condition as illustrated for the 200 ms MHT, multi-burst condition (Fig. 3C). For this condition, the number of sound bursts played during the actual hold time was determined for each individual trial across all animals (e.g., Fig. 3C, x-axis, red dots). Using a linear piece-wise regression, we fit this data (Fig. 3C, black line) and estimated the proportion of sound bursts at the median hold time (Fig. 3C, black dot), as shown for this example condition (Fig. 3C). The proportion of sound bursts heard was estimated for all median hold times and sound conditions (Fig. 3E) using the same approach.

**Figure 3:**
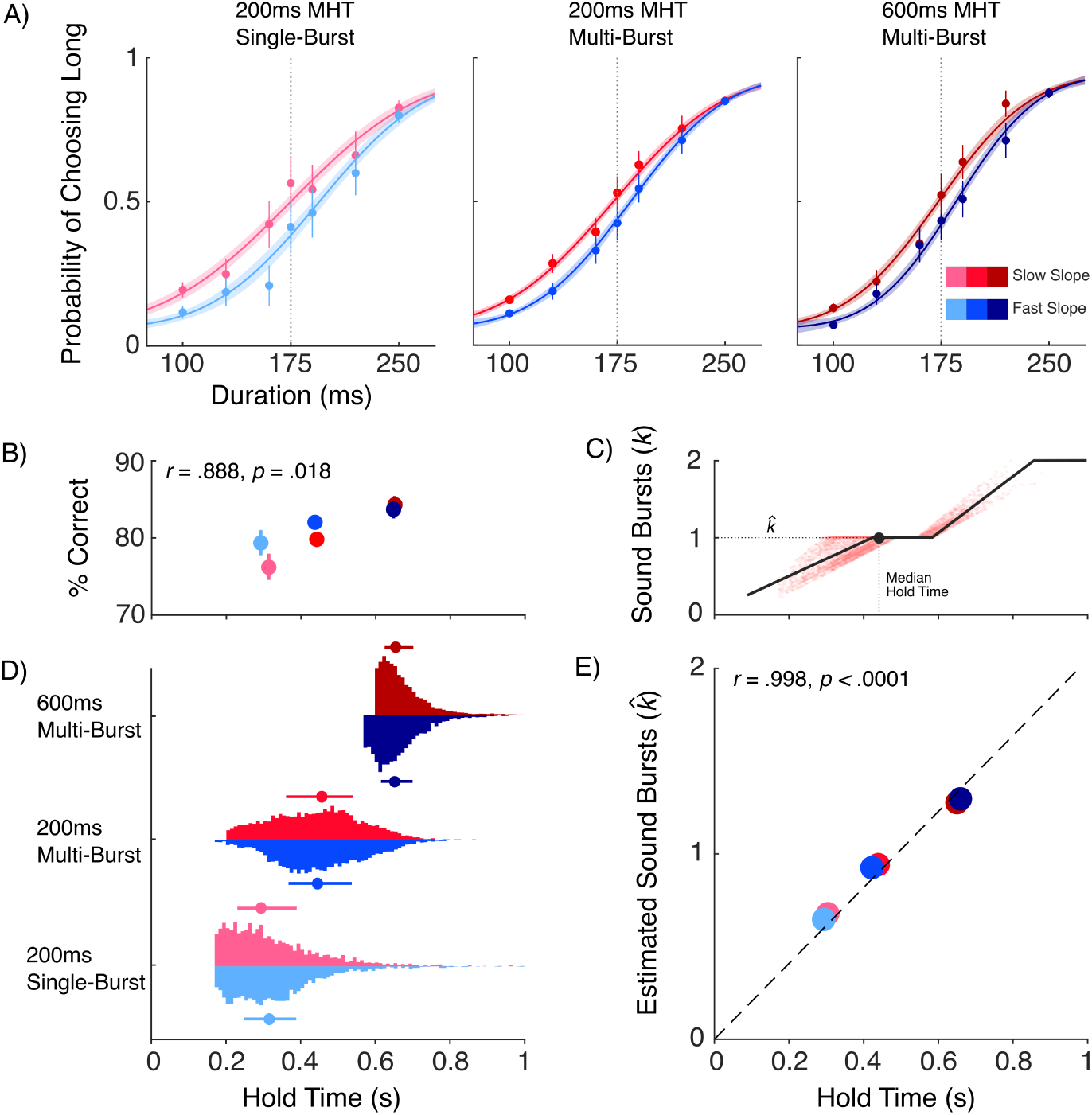
Sound onset-offset slopes and the number of sound bursts heard both impact perceptual judgment of sound duration. A) Judgment of sound duration varies as a function of sound onset-offset slope (slow versus fast, blue versus red) and MHT (left, middle and right panels). The mean choice probability (filled circles) and S.E.M. (verticle bars) for judging sound duration as “long” are plotted for each of the seven sound durations tested. To quantify the effect of sound onset-offset slope on sound duration judgment, maximum likelihood psychometric fits are generated for all MHT and sound slope conditions. Lighter colors indicate less evidence accumulation (correspondent with the MHT condition). Translucent bands indicate 95% confidence intervals (non-parametric bootstrap iterations = 400). B) Illustration of the estimated number of bursts 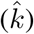 heard during a given trial with respect to the actual median hold time (based on piecewise regression, METHODS and supplementary material). C) Percent correct (%Correct) sound duration judgment increases as the MHT is increased and the two are positively correlated (r = 0.888, p = 0.018). The mean and S.E.M. of the percent correct responses are plotted as a function of the MHT (x-axis) for all six task and sound conditions. D) Violin histograms of the probability distribution of actual hold times for every trial (total trials= 22,756). E) The proportion and number of sound bursts heard increases with the actual median hold time (see supplementary material). Data shows a near absolute correlation (*r* ≈ 1).

### Descriptive model of psychometric function

To quantify how sound onset-offset slope and task conditions impact sound duration judgement, choice response data were fit with a standard modified sigmoid function using the Palamedes toolbox (Prins and Kingdom, 2018). The generic form of this psychometric function is a sigmoid link function scaled to lie between asymptotic lapse rates:

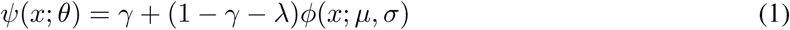

Where *θ* denotes the set of parameters {*μ, σ, γ, λ*}, and *ϕ* denotes the cumulative normal function. For equation 1, *x* refers to the sound plateau duration, *ψ* refers to probability a choice classifying the sound as long, *σ* corresponds to the standard deviation of the cumulative normal function. The standard deviation, *σ*, is linearly and inversely related to the sigmoid function slope. Thus, the sigma (*σ*) parameter and slope quantify perceptual variance and sensitivity, respectively, for judging short versus long sound durations. Accordingly, the sigma parameter indexes “sensory noise” or variation. The μ parameter corresponds to the mean of the cumulative normal distribution of choice probabilities. If we assume uniform priors and equal rewards for long versus short durations, *μ* defines the *x*-intercept and the point of subjective equality (*p*_*x*_ = 0.5), also known as the bias point, on the response function.

Rather than fitting all conditions (2 slope conditions x 3 hold time conditions) with independent sets of 4 parameters each, we sought to find the most constrained descriptive model that accounted for the data. We did this using Palamedes’ model comparison feature, which only allows parameters to vary between conditions if warranted by a model comparison (transformed likelihood ratio test), and constrains them to be fixed across conditions otherwise. (Prins and Kingdom, 2018)

### Normative (Bayesian decision-theoretic) model incorporating natural sound statistics

To examine whether natural co-variations in onset-offset slopes and durations can predict how onset-offset cues impact sound duration judgement, we constructed a Bayesian decision-theoretic model of sound epoch duration judgements.

Let the true onset-offset slope of a sound be 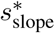 and its duration be 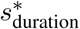 We assume that animals maintain a prior belief about the joint occurrence of these features *p*(*s*_duration_, *s*_slope_) based on natural statistics. We estimated this “natural statistics prior” based on the joint probability distribution of onset-offset slopes and durations found in natural alarm vocalizations, approximated by a bivariate Gaussian density (Fig. 1). We tested three different possibilities - that animals’ duration decisions were affected by only the onset slope, only the offset slope, or the joint distribution of onset and offset slopes - these correspond to different assumptions about which slope dimensions are predictive of natural vocalizations, and hence attended to. We assume that animals make duration judgements in accordance to Bayesian decision theory, by combining this prior with noisy sound duration evidence, and picking decisions that maximize expected utility, as follows:

We assume that noisy duration observations on any given trial *x*_duration_ is drawn from a Gaussian centered around the true duration, with a standard deviation of *σ*_*s*_.

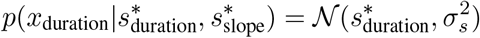

We allow for the possibility of different levels of noise depending on the onset-offset slope: 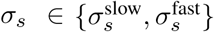 Further, we assume that the true onset-offset slope on the trial is known: 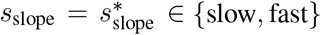 since “fast” and “slow” slopes were chosen to be at extreme ends of rats’auditory neuron slope response fields and discrimination performance, assessed previously (Lee et al., 2016; Osman et al., 2018).

Then the likelihood of the hypothesized duration *s*_duration_ on a given trial is a Gaussian function, centered around the observation *x*_duration_

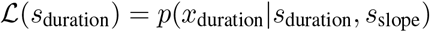

We assume that rather than receiving just 1 observation, animals receive a constant rate of independent observations over time i.e. *x*_duration_ = {*x*_1_, *x*_2_, …, *x*_*k*_}. Assuming that animals integrate these optimally, this yields a total likelihood that is the product of likelihoods for each individual observation, hence reducing in width as an inverse function of the number (i.e., Fig. 3B the number of observed sound bursts, 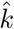) or the proportion (*k*) of observations in a given trial. We assume the proportion of observations (*k*) to be a linear function of the hold time in a given trial (Fig. 3B and 3E): *k* ∝ *t*_hold_,

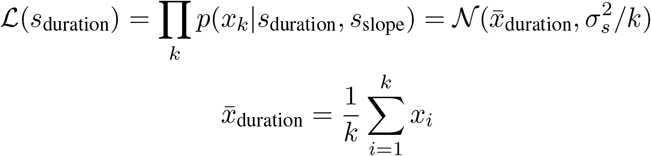

Let the conditional prior *p*(*s*_duration_|*s*_slope_) evaluated at the current trial’s slope have a mean of *μ*_prior_ and a standard deviation of *σ*_prior_. Then the posterior belief about duration, for a given set of noisy observations and slope condition is given by Bayes rule:

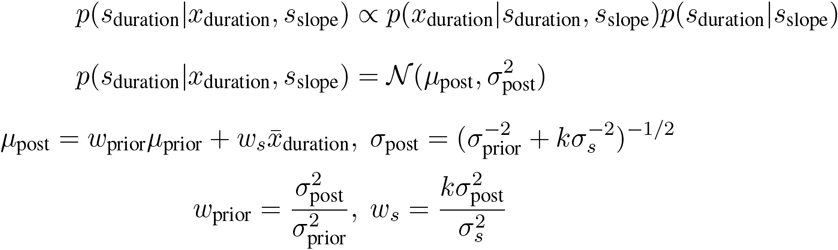

Note that as the animal receives more observations, the influence of the stimulus increases and the influence of the prior reduces.

The probability of a “long” duration is given by the integral of the posterior density beyond the true category boundary *μ*_0_, which we assume to be known. The maximum utility decision rule, assuming knowledge of rewards and priors, involves deterministically choosing “long” judgements when the posterior mean exceeds the category boundary, and “short” otherwise.

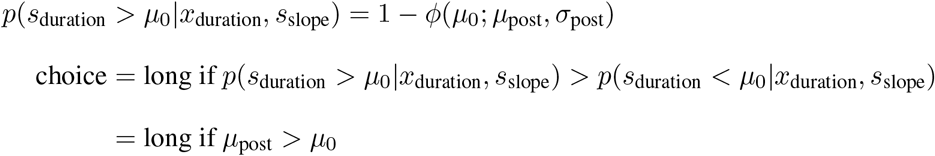

However, since the posterior mean depends on noisy observations, the probability of choosing a “long” judgement for a given true duration requires marginalizing over possible noisy values of *x*_duration_(eq. 7, Ma (2019))

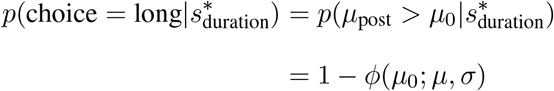

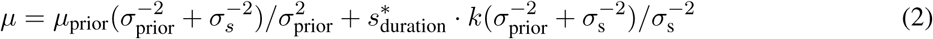

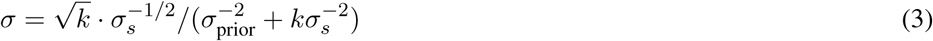

This yields a cumulative normal psychometric function (eq. 6) with midpoint *μ* and inverse slope σ. We augment this psychometric function with lapse rates, assuming that animals occasionally lapse due to fixed motor errors or inattention (Pisupati et al., 2021).

In the inattention model, *p*_lapse_ is the probability of not attending, and *p*_guess_ is assumed to be proportional to the prior probabilities of each category i.e. *p*_guess_ = *p*(*s*_duration_ *> μ*_0_|*s*_slope_), while in the motor error model *p*_lapse_ is the probability of motor error, and *p*_guess_ is assumed to be 0.5

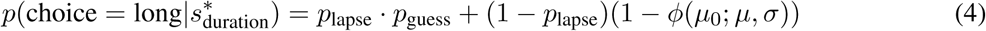

This yields a psychometric function that can be fit to the behavior in any given condition, with 2 free parameters: *σ*_*s*_ and *p*_lapse_ for both “inattention” and “motor error” lapse models. Across conditions, we force *p*_lapse_ to be the same, based on a preliminary analysis that showed empirical lapse rates to be the same across conditions. The sensory noise parameter *σ*_*s*_ is either fixed across conditions (“fixed” noise models) or allowed to vary between different slope conditions (“variable” noise models). We fix *k* = 1 for the single burst 200ms hold time condition, allowing the *k* on other conditions to reflect relative integration times (fig. 3). For the “perfect integration” model, *k* is fixed to be proportional to the empirical hold times, yielding no extra free parameters. We also entertain an “imperfect integration” model, that allows *t*_integration_ < *t*_hold_, yielding 2 extra free parameters.

For each of the three prior types (onset, offset, and both), 8 model variants (2 lapse models × 2 noise models × 2 integration models) were fit using maximum likelihood fitting, using MATLAB’s fmincon function. Thus, all together we compared 24 different models (3 prior types × 8 model variants). Model comparison was performed using Bayesian Information Criterion (BIC) and Akaike Information Criterion (AIC) as detailed (figure 5B, supplementary table 2). We also fit synthetic data generated from each model and performed a similar model comparison, in order to assess model recovery.

## RESULTS

It is known that the slope of a sound’s onset can dramatically change how we perceive its duration (Stecker and Hafter, 2000; Grassi and Darwin, 2006; Friedrich and Heil, 2017) and yet the basis for this perceptual interaction remains unknown. Here, we propose such cue interactions may reflect expectations based on prior experience hearing co-variations in the onset slope and durations of natural sounds. To examine this, we first quantify the statistical distributions of these cues in recordings of natural rodent alarm vocalization sequences (Methods, Fig. 1). As the name implies, alarm vocalizations are used to communicate an alarm to other rodents when they are distressed or alternately when they are defeated during rough and tumble play (Wöhr and Schwarting, 2008; Thomas et al., 1983; Saito et al., 2019). When rodents hear synthetic versions of these alarm vocalizations, they display stereotyped social responses provided the proper combinations of pitch and temporal cues are incorporated (Wöhr and Schwarting, 2008; Saito et al., 2019). Confirming prior studies, natural rodent alarm vocalizations examined here have a pitch or fundamental frequency around 22 kHz (Fig. 1A1, A2) and durations ranging from 50 to 1500 ms (Fig. 1B, C). In addition to these known acoustic features, we find the vocalizations with the shortest durations are statistically more likely to have faster onset slopes (Fig. 1C1). Conversely, the vocalizations with the longest duration are statistically more likely to have slower onset slopes (Fig. 1C1). Accordingly, the onset slope is inversely correlated with vocalization duration (Fig. 1C1, Onset slope: r=-.383 and p = 1e-47). In contrast, vocalization offset slopes have a weak positive correlation with duration (Fig. 1C2, Offset slope: r=.125 and p = 5e-6). These correlations between onset, offset and duration extend the list of known timing cues that identify vocalization type (Khatami et al., 2018; Saito et al., 2019).

Given the statistical co-variations reported here, we hypothesize that the slope of a sound would strongly influence the perceptual judgment of its duration. To test our hypothesis, we create synthetic vocalizations or sound bursts with plateau duration (Methods) ranging from 100 to 250 ms, which falls within the range of natural alarm vocalizations (Fig. 1C). Our sound design allows us to symmetrically vary the sound onset-offset slope independent of the sound duration (Methods). Thus, we are able to generate synthetic vocalizations with a range of durations (100-150 ms) that all have the same symmetric onset and offset slopes (Fig. 1B, bottom panel). To test the effects of the slope cue on duration perception, we generate one set of synthetic vocalizations with slow onset-offset slopes and another set with fast onset-offset slopes (Fig. 1E, red versus blue, respectively). Animals were trained initially on sound sequences with slow onset-offset slopes until they reach a high-performance criterion (Methods). Importantly, the normalized slow onset-offset slope used here is 84 A/s (Fig. 1E), which falls within the range we observe in the natural alarm vocalizations with plateau durations ranging from 100 to 150 ms (Methods, Fig. 1C). In contrast, the fast onset-offset slope used here is about two-fold faster than the fastest onset slope observed in the natural vocalizations but still within the range of slopes evoking significant neural responses in rats (Lee et al., 2016). Multivariate Gaussian fits of the joint distribution of natural vocalization slope and durations (Fig. 1C1, C2) are used to quantify the co-occurrence of sound slope and duration cues (Methods, Fig. D1, D2). In theory, prior experience hearing natural sounds such as alarm vocalizations and the statistical covariations therein could influence animals’ inferences about sound duration. Moreover, co-variation of onset or offset slopes alone or together could be used to infer duration. Accordingly, these three cue-combination scenarios quantified with Gaussian distributions corresponding to fast and slow slopes are used to define the 1-dimensional priors over sound duration for the fast and slow slope conditions, respectively (Fig. 1F, blue versus red, respectively, Methods).

Previous studies find sound judgment can improve with an accumulation of sensory information across time and repetitions of acoustic events (Brunton et al., 2013; Liu et al., 2015; Raposo et al., 2012). This principle applies to sound duration perception (Raposo et al., 2012), though most other prior studies have probed duration perception of single sound bursts instead of sound burst sequences (Friedrich and Heil, 2017; Kelly et al., 2006). Here, we use a binomial choice task (Fig. 2) to test how animals judge the duration of synthetic sounds when onset-offset slopes are fast versus slow (Fig. 2A). The duration judgment is examined across three task conditions (Fig. 2B) using a reinforcement strategy (Fig. 2C) to train animals to perform this bimodal two-alternative forced-choice behavioral task (Fig. 2D). Our three task contingencies allow us to examine sound duration judgment as a function of the proportional number of sound bursts heard (Fig. 3). Though the required minimum hold time (MHT) is fixed in the three task contingencies, animals may hold longer than the MHT and the actual hold time varies on a trial-by-trial basis (Methods, Fig. 3D, solid red and blue dots). For example, when MHT is 200 ms and multiple sound bursts are played, the actual hold time is 445 ms (Fig. 3C, vertical dotted line). Under the latter condition, animals hear on average 1 single sound burst before releasing from the center port (Fig. 3C, horizontal dotted line). Conversely, when MHT is 600 ms and the actual median hold time is 652 ms (Fig. 3D, red dot, top histogram), the number of sound bursts heard is 1.28 on average. Thus, the proportional number of sound bursts heard increases when MHT is increased from 200 to 600 ms and multiple sound bursts are heard (Fig. 3E). In the third task contingency where the MHT is 200 ms and only a single sound burst is ever played, animals typically heard only a fraction (0.74) of one sound burst before making a duration judgment (Fig. 3E). In all task conditions, animals judge 7 different sound durations as short versus long (Fig. 2C) based on reward and time-out contingencies (Fig. 2D, Methods). Animals initially learn to hold for a minimum of 600 ms while hearing a sequence of synthetic vocalizations with a varied but average repetition rate of 2 Hz (Fig. 2B, 600 ms MHT, multi-burst). After reaching the task performance criteria (Methods), duration judgment is tested for sounds with slow versus fast onset-offset slopes in alternating test blocks (Fig. 2A). Animals then progressively learn to perform the sound duration judgment task while holding for a minimum of 200 ms and hearing only one synthetic vocalization (Fig. 2B).

Though prior studies have demonstrated that sound slope cues alter duration perception none have examined how accumulating more sensory evidence impacts this cue interaction. Here, we find that sound duration judgment is significantly impacted by both the sound slope cues and by hearing proportionally more sound bursts (Figs. 3 and 4). Population performance for judging seven different sound durations is quantified as the mean choice probability for judging sounds as long in duration (Fig. 3A, filled circles, Methods). Initially, behavioral responses are fit with a standard sigmoidal response function (Methods, Fig. 3A, red and blue lines) in order to quantify performance metrics (Fig. 4). For all three MHT and task conditions, there is a rightward shift in the perceptual boundary or ”bias” (Fig. 3A, vertical dotted lines) for sound duration judgment when sounds have fast (Fig. 3A, blue lines) versus slow (Fig. 3A, red lines) onset-offset slope. This rightward bias indicates that animals are judging all sound durations as shorter when their onset-offset slope is faster. This perceptual bias is most pronounced for the fast slope condition, and when animals haven’t accumulated much sensory information i.e. when they hear a maximum of one single sound (Fig. 3A, left panel) and on average only a fraction of a single sound burst (Fig. 3E, light blue, and pink dots). This perceptual effect is readily appreciated by comparing the bias parameter when hearing a single burst with slow (Fig. 4A, SB) versus fast (Fig. 4B, SB) onset-offset slopes. Independent of the sound onset-offset slope, sound duration judgment becomes sharper, and psychometric response functions steeper, as more sound bursts are heard (Fig. 3A, 600 ms vs 200 ms MHT, right versus left panels). Accordingly, there is a rank order decrease in the inverse sensitivity (Fig. 4C, D) which corresponds to an increase in judgment accuracy across task conditions where animals hear proportionally more sound bursts. This effect is observed for sounds with slow or fast onset-offset slopes suggesting that it is primarily related to the accumulation of sensory information and not sound slope. Finally, changing the sound onset-offset slope or number of sound bursts heard has no impact on the overall performance levels and relative performance lapse for judging the shortest and longest duration sounds (Fig. 4E, F). Together, these results indicate that sound slope cues can bias duration perception and that accumulating more sensory evidence can effectively reduce this bias.

**Figure 4:**
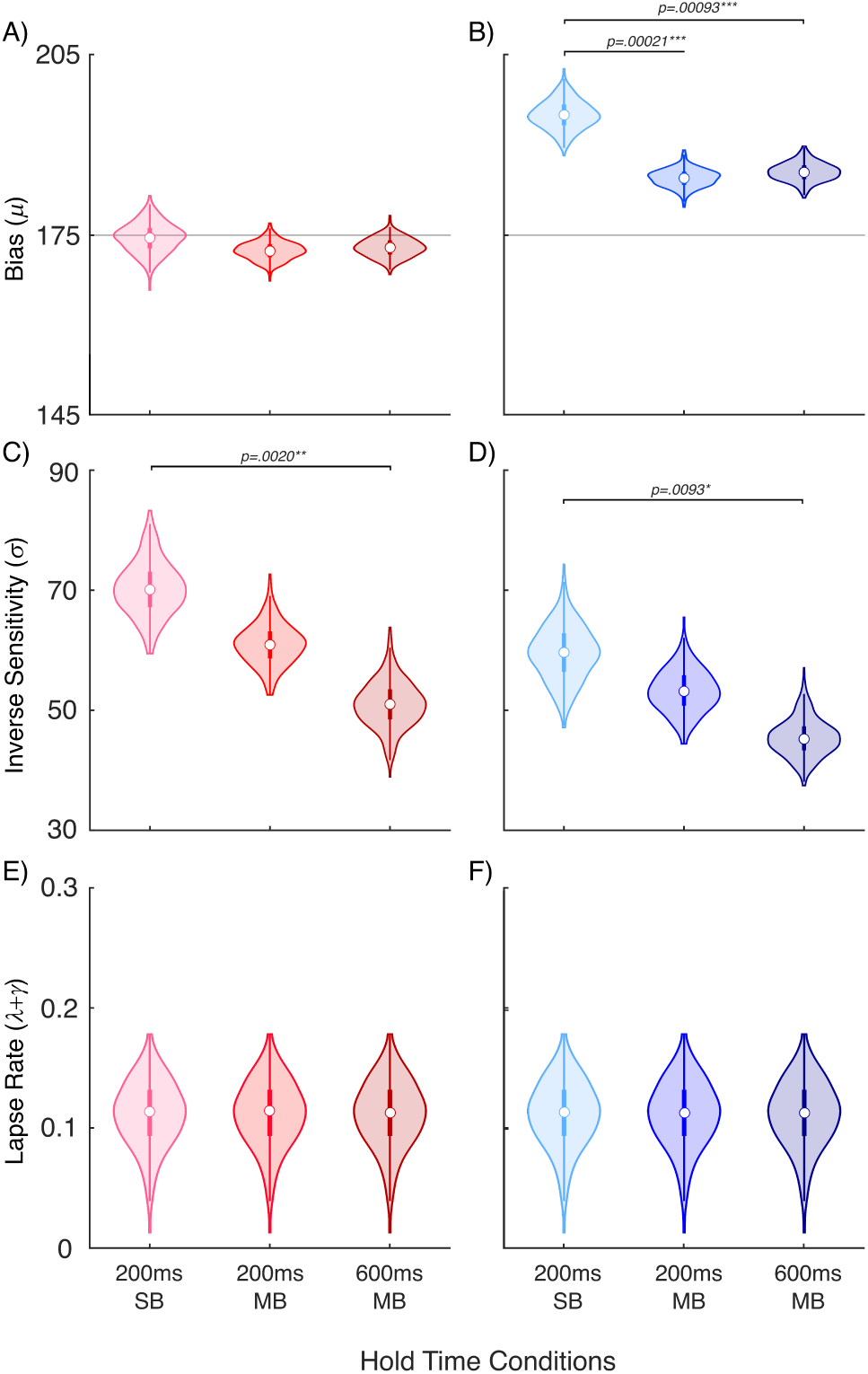
Distinct shifts in perception with change in onset-offset slopes versus change in number of bursts heard. Psychometric response function bias (*µ*), inverse sensitivity (*σ*) and lapse parameters (*γ*+*λ*) sampling distributions are shown for all task conditions (non-parametric bootstrap iterations = 400). A-B) The bias is shifted to higher values under all three MHT conditions when sound onset-offset slopes change from slow to fast (two-tailed z-test - 200ms SB: p=9.5e-8, z=5.77; 200ms MB: p=1.4e-7, z=5.71; 600ms MB: p=2.2e-7, z=5.62). C-D). Inverse sensitivity or *σ* shows a consistent decrease with respect to the hold time condition (z-test p-values shown in figure). E-D) Lapse rates are fixed across hold time and and sound slope condition (mean [+/- SEM]= .113 [+/-.028]). * p*<*.05,** p*<*.01, ***p*<*.001

Although our behavioral results indicate that sound slope cues impact sound duration judgments, the underlying principles driving this temporal cue and task interaction remain unclear. To examine whether such interactions stem from expectations based on prior experience hearing natural sounds,we fit the behavioral responses using a normative, Bayesian decision-theoretic model of decision-making (Fig. 5). As illustrated in the joint probability distributions of durations and slopes, onset and offset slopes both co-vary with the duration of natural alarm vocalizations (Fig. 1). We utilized the Gaussian fits of these empirical joint distributions as parameter-free, “natural statistics priors” to infer duration in our Bayesian model (Fig. 1E). To fit the data, we compare three different prior types (Fig. 5B, table with rows reflecting prior type) derived from three different natural statistical joint distributions for duration and onset slope (onset, Fig. 1F, top), duration and offset slope (offset, Fig. 1F, middle) or duration with both onset and offset slopes (both, Fig. 1F, bottom). The model had free parameters to account for three potential sources of errors in decisions: “noise” in sensory observations, “lapses” or random decisions (Pisupati et al., 2021), and suboptimal “integration” strategies. Accounting for these three sources of noise in our models with different integration constraints generates a total of 8 model variations tested (Methods, Fig. 5B, table with three types of column subdivisions: lapse type, noise constraint, and integration type). The noise in duration observations could either be the same across slope conditions (“fixed”, 1 parameter across conditions) or differ for different slope conditions (“variable”, 2 parameters for the two slope conditions). Lapses, or decisions made irrespective of duration evidence, could arise due to motor errors and hence be made randomly (choosing either decision with a probability of 0.5), and occur with a fixed probability across conditions (“motor error”, 1 parameter across conditions), or arise from inattention and hence be made with a bias reflecting that condition’s prior, and occur with a variable probability across conditions, (“inattention”, 2 parameters for the two slope conditions). The integration of evidence across multiple bursts was either perfectly optimal, with the number of effective bursts being fixed to be equal to the empirical hold times (“perfect”, no extra free parameters), or suboptimal, with the effective number of bursts being less than the empirical hold times (“imperfect”, with 2 additional free parameters). The different assumptions about these three sources of errors, combined with the three different prior types gave rise to a total of 24 combinations, which we compared using factorial model comparison (Ma, 2019).

**Figure 5:**
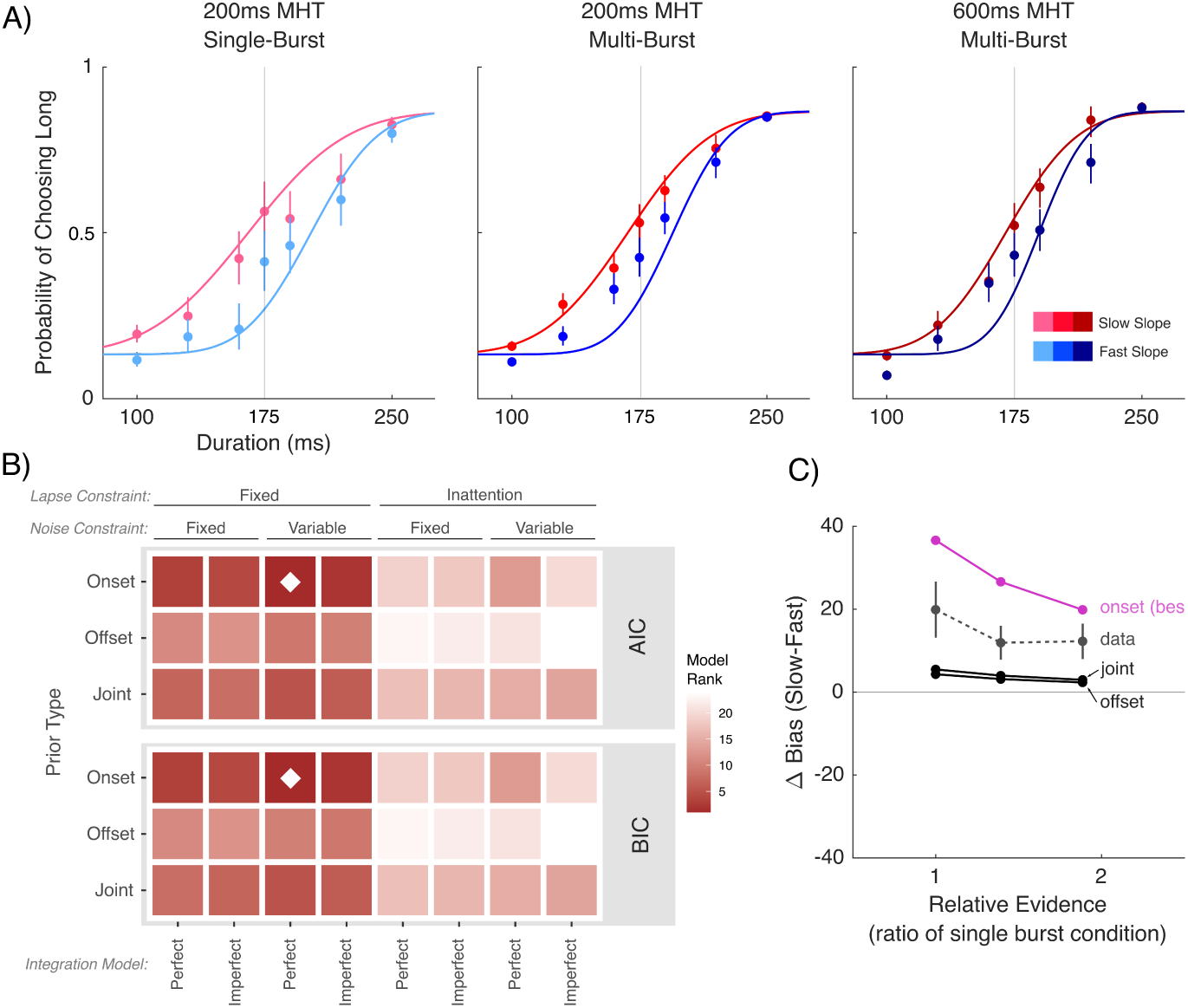
Bayesian decision theoretic (BDT) model of observed bias. A) Model fits from the best-fitting BDT model (solid lines) overlaid onto raw decision probabilities (points) from the behavioral experiment. B) Factorial model comparison of each of the three types of parameter constraints (lapse, noise and integration) for the three types of prior (onset, offset and both). Models are rank-ordered so that the model with the lowest information criterion (Akaike and Bayesian) is 1 (the best model, darkest color) and the one with the highest information criterion is 24 (the worst model, lightest color). Diamond denotes the best model.

When we fit the behavioral data with the twenty-four different models, we find the overall best-fitting model according to Bayesian information criterion (BIC) and Akaike information criteria (AIC) used the empirical prior based on sound onset slopes and durations, and required only three free parameters - variable sensory noise parameters across slope conditions, and a fixed lapse probability due to motor errors across conditions, with perfect integration of evidence determined by the empirical hold times (Fig. 5A, 5B, diamond). In contrast, across all model variations, priors based on the offset slope alone or based on a combination of onset and offset slope did not accurately predict shifts in sound duration judgment with sound slope (Fig. 5B, offset and both). Additionally, sensory noise but not lapse probabilities must vary to account for response differences across behavioral task conditions. Importantly, our model is highly constrained, using an entirely parameter-free empirical prior to account for perceptual biases, and only 3 additional free parameters to account for other sources of errors. In comparison, the standard sigmoidal model requires thirteen free parameters (with a total of 6 individual bias parameters) to accurately fit the response data (Fig. 3A). Hence, incorporating natural sound statistics offers a parsimonious explanation for the perceptual misjudgements in sound duration caused by varying sound slopes. Finally, our Bayesian model captures the inverse relationship observed between the empirical hold times and the perceptual bias as well as inverse sensitivity, since integrating more sensory evidence overcomes the influence of the prior, leading to less biased and more accurate decisions (Fig. 5C). Since the best fitting model performed perfect integration, it required no additional free parameters to capture this effect, directly using the empirical hold times as a parameter-free proxy for the number of accumulated evidence bursts. In summary, model fits indicate that two key factors account for (mis)judgments of sound duration - prior experiences with co-variations in natural sound statistics, and the amount of accumulated sensory evidence. Together, these results offer a principled and normative explanation for why and how one auditory temporal cue can bias the perception of another, and how accumulating sensory information can overcome these biases.

## DISCUSSION

Previous studies find the rate of sound onset dramatically influences the perception of sound duration but the underlying principles for these cue interactions remain unknown (Cumming et al., 2015; Paquette and Peretz, 1997; Stecker and Hafter, 2000; Grassi and Darwin, 2006; Friedrich and Heil, 2017; Bizley and Cohen, 2013). We previously demonstrated that natural co-variations between duration and other temporal cues can be used to differentiate vocalization type across many animals including humans (Khatami et al., 2018). Using a similar approach here, we find a strong inverse correlation between the distribution of onset slopes and durations of rodent alarm vocalizations (Fig 1). Accordingly, vocalizations with faster onset slopes are more likely to be short in duration. Given this correlation, onset slopes could serve as a predictive cue for vocalization duration. In contrast, offset slope and duration only have a weak positive correlation. These observations lead us to hypothesize that perception of sound duration should be biased by sound onset slopes, more so than offset slopes. Behaviorally, we find that rodents are perceptually biased to judge synthetic vocalizations with fast onset-offset slopes as being shorter in duration (Fig. 3, 4). To gain insight into this “mis-judgment” of duration and explore the potential contributions of onset and offset slope statistics, we model the behavior with a normative, Bayesian decision-theoretic model. We find that the behavioral data is best fit by a model that incorporates the joint statistics of durations and onset slopes of natural vocalizations as a prior (Fig. 5A, blue curves; Fig. 5B, diamond). This supports our hypothesis that onset slope more strongly biases the perception of duration than offset slope due to its natural co-variations with duration, and accounts for the behavioral biases observed in the present study. Since our model accounts for these biases by using empirical priors derived from natural vocalizations, and empirical hold times as a proxy of accumulated evidence, it requires far fewer parameters than standard psychometric functions to capture the observed biases (Fig. 3). Models that incorporate co-variations in onset slope and duration perform better than similarly constrained models incorporating co-variations of offset and duration, or the combined co-variations of onset and offset slopes with duration (Fig. 5B, offset and both). Moreover, our Bayesian model captures the behaviorally observed decrease in bias (Fig. 5C) and improvement in sound duration sensitivity (Fig. 3B) as animals listen to and integrate more sensory information across multiple vocalizations, reducing the influence of the prior. In summary, our results demonstrate that prior experience with the natural co-variations in onset, offset and duration cues can explain why onset slope cues heavily bias perception of sound duration, with this perceptual bias reducing if perceptual evidence is integrated over longer time windows.

Cue integration both within and across sensory modalities has been shown to follow principles of Bayesian inference in a number of studies in humans (Trommershauser et al., 2011) as well as rodents (Raposo et al., 2012; Madl et al., 2014; Nikbakht et al., 2018; Sheppard et al., 2013). According to these principles, animals integrate information from multiple cues if they expect them to arise from a common source that produces correlated measurements across cues. Such expectations can be formalized as a “coupling prior” between cues, reflecting statistical regularities learned with prior experience (Spence, 2011). Such priors are beneficial (i.e. lead to improved accuracy) in natural environments and tasks that respect these correlations, and especially beneficial when sensory information about one or more cues is limited or noisy. However, the same priors can be detrimental (i.e. lead to biases and impaired accuracy) in tasks that do not respect these natural, learned regularities. Accordingly, in the present study when the synthetic vocalization duration is the task-relevant cue and the onset-offset slopes are artificially fast compared to natural statistics, duration judgment is biased and performance drops. In a similar vein, rats and mice performing a visual rate-discrimination task (Odoemene et al., 2018) are influenced not just by the rewarded relevant variable (i.e. event rate) but also by the total event count. While this may be beneficial on most trials when rate and count are correlated, it can lead to incorrectly biased decisions on “catch” trials when the two are varied independently. These findings support the idea that there are advantages and disadvantages to relying on environmental priors, especially when generalizing them to new environments, and decision-making systems will need to flexibly tune how much they generalize in order to remain adaptive.

How can one be sure that biases observed in a given task are the result of biased priors, rather than other biasing influences on decisions? Unconstrained Bayesian models might be overly flexible and capable of accounting for a vast range of erroneous behaviors through the use of mismatched priors, and hence difficult to falsify (Rahnev and Denison, 2018). This is why we instead opt for the approach of constraining the parameters of the prior in our model entirely based on empirical natural statistics, with the sole assumption being that animals’ judgments about synthetic vocalizations are biased by their prior beliefs about sounds such as vocalizations. Moreover, decisions in Bayesian models are made by combining priors with incoming samples of sensory evidence, each weighted by their respective certainty. Hence prolonged sampling and integration of sensory evidence can lead to more accurate decisions by offering more certain evidence and correcting for any biases from the prior, a feature evident in the behavioral data and captured by our model based on empirical sampling times. These results extend previous work in support of the ability to optimally accumulate auditory information to improve perceptual accuracy (Brunton et al., 2013)

What neural substrates could underlie these Bayesian computations? In order to maximize efficiency, neurons in the brain should utilize codes that match the statistics of the signals they represent (Gervain and Geffen, 2019; Carruthers et al., 2013). Accordingly, theoretical work on efficient coding has proposed that prior distributions could be implicitly represented in the distribution of tuning curves in neural populations, with regions of higher prior probability being tiled more densely by more selective neurons (Ganguli and Simoncelli, 2014a). Consequently, encoding a correlated “coupling prior” across multiple cues would entail joint (rather than independent) encoding of these cues, with sharper and denser tuning to cue combinations congruent with the prior (Yerxa et al., 2020a). Such joint encoding schemes would encourage integration, prioritizing the efficient inference of common sources over the accurate reconstruction of component cues (Zhang et al., 2019). At the same time, independent encoding schemes would remain advantageous when no common source is detected and integration is not warranted (Zhang et al., 2019).

The simultaneously parallel and hierarchical structure of cortical pathways offers a possible candidate for a flexible Bayesian inference (Rohe et al., 2019). Following a general rule observed in other sensory cortices, as one transitions from primary to secondary auditory cortices, neurons respond to dynamically changing sensory stimuli on increasingly longer timescales (Hamilton et al., 2018; Wang and Kennedy, 2016; Chaudhuri et al., 2015; Lee et al., 2016; Johnson et al., 2020). We and others have shown previously that primary and secondary auditory cortices encode multiple temporal cues including onset and offset timing, duration, and rhythmicity cues in sound sequences (Lee et al., 2016; Read and Reyes, 2018). However, primary cortical neurons respond to and encode sound onset-offset slope and sound rhythmicity independently (Lee et al., 2016), and more accurately categorize the sound’s onset-offset slope and rhythmicity than those in secondary auditory cortices (Osman et al., 2018). Likewise, primary auditory cortical neurons accurately encode variations in spectral and temporal cues in natural vocalization sequences (Lee et al., 2016; Storace et al., 2011; Carruthers et al., 2013; Gervain and Geffen, 2019). In contrast, secondary auditory cortical neurons respond to and encode these cues in a joint manner. For example, secondary auditory cortical neurons that respond preferentially to sounds with slow onset-offset slopes tend to have more sustained spike-timing responses and consequently a slower repetition rate or rhythmicity sensitivity (Lee et al., 2016). This co-variation in neural sensitivity to the two temporal cues can be used to objectively differentiate sound sequences (Osman et al., 2018). Thus, its neural spiking patterns, much like the natural sensory statistics themselves, can provide temporal cues that distinguish natural vocalizations (Khatami et al., 2018; Elie and Theunissen, 2019; Carruthers et al., 2013). This raises the interesting possibility that the primary auditory cortical area provides a neural substrate for more accurate estimations of separate sources through independent encoding, while secondary auditory cortical areas allow for more efficient probabilistic inference of common sources. Accordingly, secondary auditory cortex might be expected to encode the joint statistical co-variations found in natural sounds such as vocalizations, providing the “top-down” neural substrate for a “coupling prior”. Though future studies are needed to establish this link, auditory cortices contain the neural code to represent multiple temporal cues and support their optimal integration with prior experience. This neural code may in turn be relayed to downstream areas such as the secondary motor cortex or striatum for further integration over time (Erlich et al., 2015; Yartsev et al., 2018), or with value (Pisupati et al., 2021) to support optimal decision-making.

## Supplemental Materials

**Supplemental Figure 1:**
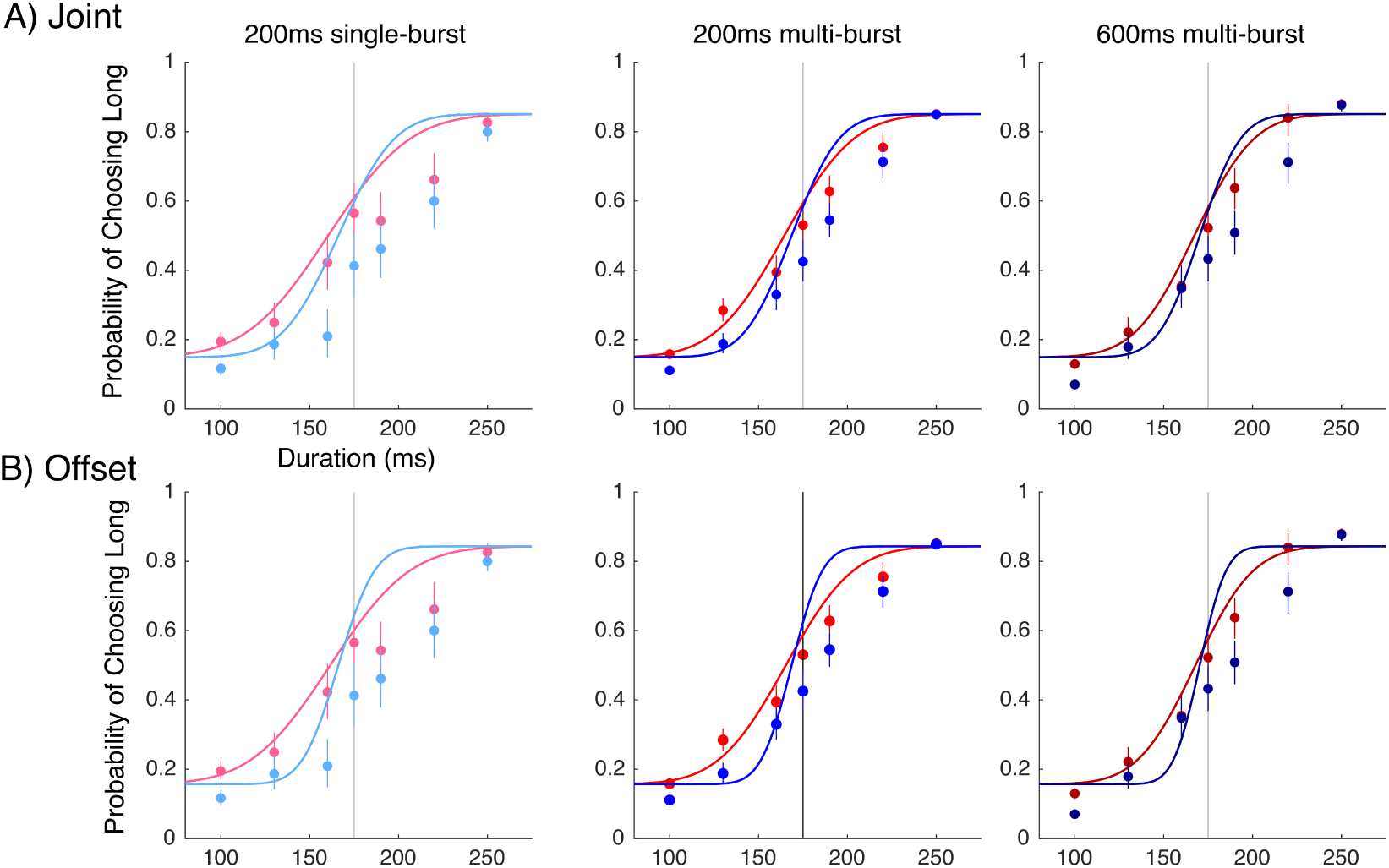
Best model fits according to AIC for offset and joint prior types.

**Supplemental Table 1:** Model fit metrics for all models compared in the study.

| Prior Type | Noise Constraint | Integration Model | Lapse Constraint | $\Delta$ BIC | $\Delta$ AIC | nLL |
| --- | --- | --- | --- | --- | --- | --- |
| onset | fixed | perfect | fixed | 76.37 | 84.59 | 12834.83 |
| onset | fixed | perfect | inattention | 7271.68 | 7279.89 | 16432.48 |
| onset | fixed | imperfect | fixed | 95.55 | 87.33 | 12834.2 |
| onset | fixed | imperfect | inattention | 7220.65 | 7212.44 | 16396.75 |
| onset | variable | perfect | fixed | 0 | 0 | 12791.53 |
| onset | variable | perfect | inattention | 5706.65 | 5706.65 | 15644.86 |
| onset | variable | imperfect | fixed | 18.02 | 1.59 | 12790.33 |
| onset | variable | imperfect | inattention | 13243.25 | 13226.82 | 19402.94 |
| offset | fixed | perfect | fixed | 705.07 | 713.29 | 13149.18 |
| offset | fixed | perfect | inattention | 31603.35 | 31611.57 | 28598.32 |
| offset | fixed | imperfect | fixed | 854.48 | 846.27 | 13213.67 |
| offset | fixed | imperfect | inattention | 20168.25 | 20160.03 | 22870.55 |
| offset | variable | perfect | fixed | 698.15 | 698.15 | 13140.61 |
| offset | variable | perfect | inattention | 13535.87 | 13535.87 | 19559.47 |
| offset | variable | imperfect | fixed | 716.38 | 699.94 | 13139.5 |
| offset | variable | imperfect | inattention | 27684.32 | 27667.88 | 26623.47 |
| joint | fixed | perfect | fixed | 417.16 | 425.38 | 13005.22 |
| joint | fixed | perfect | inattention | 6873.15 | 6881.37 | 16233.22 |
| joint | fixed | imperfect | fixed | 431.05 | 422.83 | 13001.95 |
| joint | fixed | imperfect | inattention | 6701.02 | 6692.81 | 16136.94 |
| joint | variable | perfect | fixed | 373.03 | 373.03 | 12978.05 |
| joint | variable | perfect | inattention | 6640.6 | 6640.6 | 16111.83 |
| joint | variable | imperfect | fixed | 725.58 | 709.15 | 13144.11 |
| joint | variable | imperfect | inattention | 6486.2 | 6469.76 | 16024.41 |

**Supplemental Table 2:** Summary statistics used as the parameters for joint priors. Correlation matrix contains Pearson product-moment correlations below the diagonal and their corresponding p-values above the diagonal.

|  | Duration | Onset Slope | Offset Slope |
| --- | --- | --- | --- |
| Duration | 1 | 3.0e-39 | 2.0e-26 |
| Onset Slope | -0.38 | 1 | 4.0e-06 |
| Offset Slope | 0.12 | 0.13 | 1 |
| Mean | 586.5 | 38.5 | 10.2 |
| SD | 245.6 | 24.5 | 9.9 |

**Supplemental Figure 2:**
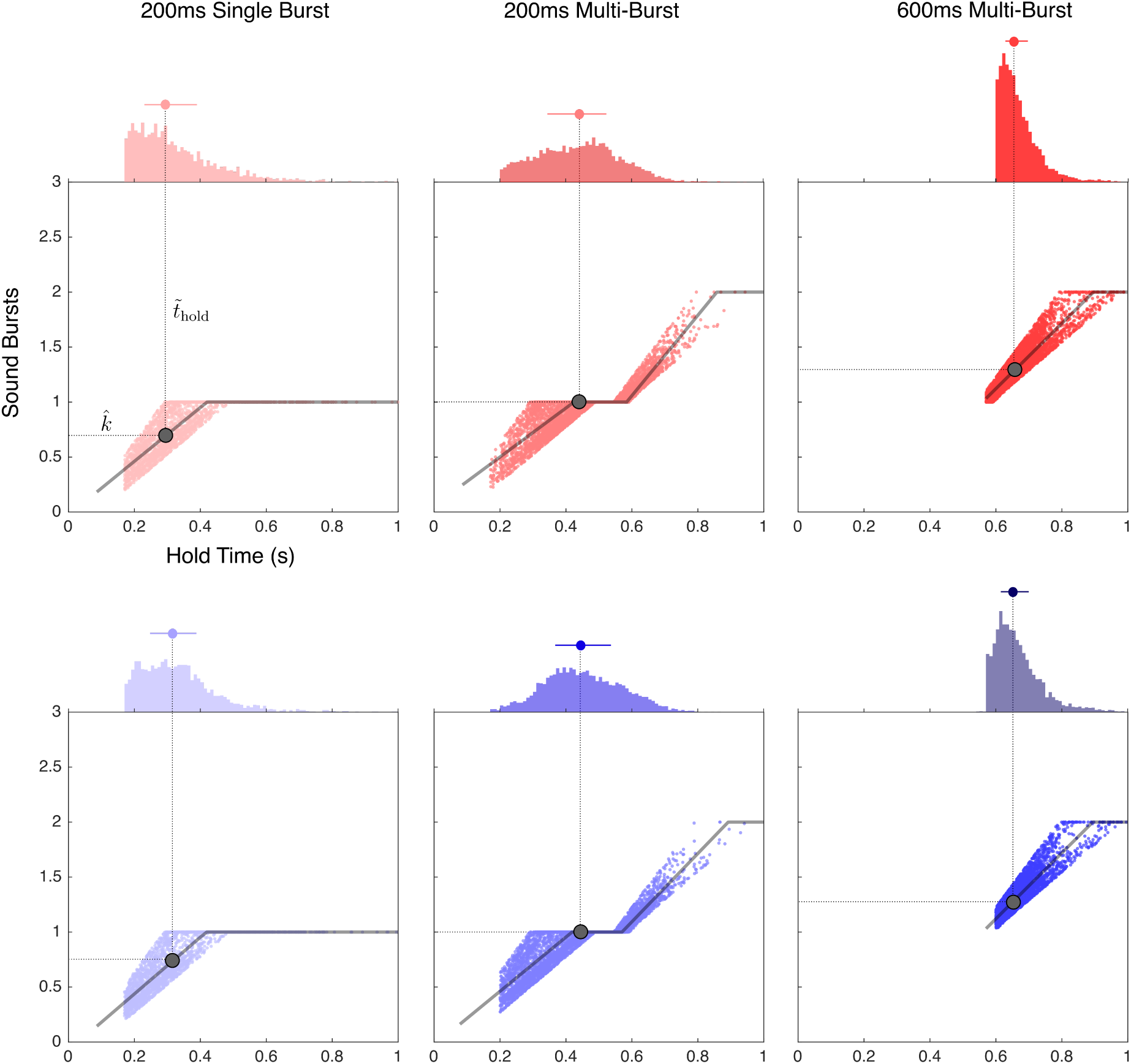
Scatterplots demonstrating the relationship between the time the animal was holding in the center port and the number of bursts heard during that time period (all trials). The residual variability is due to varying duration and the intentional jittering of the presentation of the sound stimulus. A piece-wise linear regression is fit to the data using robust least squares in order to account for heteroscedasticity. Plateaus represent the empty intervals in between sound bursts. The number of estimated bursts 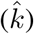 in a given condition is predicted based on the median hold time for the given condition 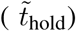. As shown in figure 3, 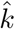 and 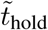 are nearly perfectly correlated (r = 0.998, p *<* 0.0001).

## Acknowledgements

H.L.R. acknowledges support from National Science Foundation NSF grant 1355065 (HLR, PI), National Institute of Health NIH DC015138 01 (HLR, Co-I), and the University of Connecticut Brain-Computer Interface Core. Grant support from the Deutsche Forschungsgemeinschaft are acknowledge fro L.M. (ME4197/2 and ME4197/3) and M.W. (WO1732/4). L.C-T acknowledges support from a CAPES Brazilian foundation grant.

## References

Belin P, Fillion-Bilodeau S, Gosselin F (2008) The Montreal Affective Voices: a validated set of nonverbal affect bursts for research on auditory affective processing. Behavior research methods 40:531–539.

Bizley JK, Cohen YE (2013) The what, where and how of auditory-object perception. Nature Reviews Neuroscience 14:693–707.

Brunton BW, Botvinick MM, Brody CD (2013) Rats and humans can optimally accumulate evidence for decision-making. Science 340:95–98.

Carruthers IM, Natan RG, Geffen MN (2013) Encoding of ultrasonic vocalizations in the auditory cortex. Journal of neurophysiology 109:1912–1927.

Chaudhuri R, Knoblauch K, Gariel MA, Kennedy H, Wang XJ (2015) A large-scale circuit mechanism for hierarchical dynamical processing in the primate cortex. Neuron 88:419–431.

Cumming R, Wilson A, Goswami U (2015) Basic auditory processing and sensitivity to prosodic structure in children with specific language impairments: a new look at a perceptual hypothesis. Frontiers in Psychology 6:972.

Elie JE, Theunissen FE (2019) Invariant neural responses for sensory categories revealed by the time-varying information for communication calls. PLoS computational biology 15:e1006698.

Elliott TM, Theunissen FE (2009) The modulation transfer function for speech intelligibility. PLoS computational biology 5:e1000302.

Erlich J, Brunton B, Duan C, Hanks T, Brody C (2015) Distinct effects of prefrontal and parietal cortex inactivations on an accumulation of evidence task in the rat. Elife 4.

Friedrich B, Heil P (2017) Onset-duration matching of acoustic stimuli revisited: conventional arithmetic vs. proposed geometric measures of accuracy and precision. Frontiers in psychology 7:2013.

Ganguli D, Simoncelli E (2014a) Efficient sensory encoding and Bayesian inference with heterogeneous neural populations. Neural Comput 26:2103–34.

Ganguli D, Simoncelli EP (2014b) Efficient sensory encoding and Bayesian inference with heterogeneous neural populations. Neural computation 26:2103–2134.

Gaucher Q, Huetz C, Gourévitch B, Laudanski J, Occelli F, Edeline JM (2013) How do auditory cortex neurons represent communication sounds? Hearing research 305:102–112.

Geffen MN, Gervain J, Werker JF, Magnasco MO (2011) Auditory perception of self-similarity in water sounds. Frontiers in integrative neuroscience 5:15.

Gervain J, Geffen MN (2019) Efficient neural coding in auditory and speech perception. Trends in neurosciences 42:56–65.

Grassi M, Darwin CJ (2006) The subjective duration of ramped and damped sounds. Perception & Psychophysics 68:1382–1392.

Hamilton LS, Edwards E, Chang EF (2018) A spatial map of onset and sustained responses to speech in the human superior temporal gyrus. Current Biology 28:1860–1871.

Jaramillo S, Zador AM (2014) Mice and rats achieve similar levels of performance in an adaptive decisionmaking task. Frontiers in systems neuroscience 8:173.

Johnson JS, Niwa M, O’Connor KN, Sutter ML (2020) Amplitude modulation encoding in the auditory cortex: comparisons between the primary and middle lateral belt regions. Journal of Neurophysiology 124:1706–1726.

Jürgens R, Fischer J, Schacht A (2018) Hot speech and exploding bombs: autonomic arousal during emotion classification of prosodic utterances and affective sounds. Frontiers in psychology 9:228.

Kelly JB, Cooke JE, Gilbride PC, Mitchell C, Zhang H (2006) Behavioral limits of auditory temporal resolution in the rat: amplitude modulation and duration discrimination. Journal of Comparative Psychology 120:98.

Khatami F, Wöhr M, Read HL, Escabí Ma (2018) Origins of scale invariance in vocalization sequences and speech. PLoS computational biology 14:e1005996.

Lausen A, Hammerschmidt K (2020) Emotion recognition and confidence ratings predicted by vocal stimulus type and prosodic parameters. Humanities and Social Sciences Communications 7:1–17.

Lee CM, Osman AF, Volgushev M, Escabí Ma, Read HL (2016) Neural spike-timing patterns vary with sound shape and periodicity in three auditory cortical fields. Journal of neurophysiology.

Liu AS, Tsunada J, Gold JI, Cohen YE (2015) Temporal integration of auditory information is invariant to temporal grouping cues. ENeuro 2.

Ma WJ (2019) Bayesian decision models: A primer. Neuron 104:164–175.

Madl T, Franklin S, Chen K, Montaldi D, Trappl R (2014) Bayesian integration of information in hippocampal place cells. PLOS one 9:e89762.

McDermott JH, Simoncelli EP (2011) Sound texture perception via statistics of the auditory periphery: evidence from sound synthesis. Neuron 71:926–940.

Melo-Thomas L, Tonelli LC, Müller CP, Wöhr M, Schwarting RK (2020) Playback of 50-khz ultrasonic vocalizations overcomes psychomotor deficits induced by sub-chronic haloperidol treatment in rats. Psychopharmacology 237:2043–2053.

Nikbakht N, Tafreshiha A, Zoccolan D, Diamond ME (2018) Supralinear and supramodal integration of visual and tactile signals in rats: psychophysics and neuronal mechanisms. Neuron 97:626–639.

Odoemene O, Pisupati S, Nguyen H, Churchland AK (2018) Visual evidence accumulation guides decisionmaking in unrestrained mice. Journal of Neuroscience 38:10143–10155.

Osman AF, Lee CM, Escabí Ma, Read HL (2018) A hierarchy of time scales for discriminating and classifying the temporal shape of sound in three auditory cortical fields. Journal of Neuroscience 38:6967–6982.

Paquette C, Peretz I (1997) Role of familiarity in auditory discrimination of musical instrument: a laterality study. Cortex 33:689–696.

Pisupati S, Chartarifsky-Lynn L, Khanal A, Churchland A (2021) Lapses in perceptual decisions reflect exploration. Elife 10.

Prins N, Kingdom FA (2018) Applying the model-comparison approach to test specific research hypotheses in psychophysical research using the palamedes toolbox. Frontiers in psychology 9:1250.

Rahnev D, Denison R (2018) Suboptimality in Perceptual Decision Making. Behav Brain Sci pp. 1–107.

Raposo D, Sheppard JP, Schrater PR, Churchland AK (2012) Multisensory decision-making in rats and humans. Journal of neuroscience 32:3726–3735.

Read HL, Reyes AD (2018) Sensing Sound Through Thalamocortical Afferent Architecture and Cortical Microcircuits In The Mammalian Auditory Pathways, pp. 169–198. Springer.

Rohe T, Ehlis A, Noppeney U (2019) The neural dynamics of hierarchical Bayesian causal inference in multisensory perception. Nat Commun 10:1907.

Roth S, Black MJ (2007) On the spatial statistics of optical flow. International Journal of Computer Vision 74:33–50.

Saito Y, Tachibana RO, Okanoya K (2019) Acoustical cues for perception of emotional vocalizations in rats. Scientific reports 9:1–9.

Sheppard J, Raposo D, Churchland A (2013) Dynamic weighting of multisensory stimuli shapes decisionmaking in rats and humans. J Vis 13.

Spence C (2011) Crossmodal correspondences: A tutorial review. Attention, Perception, & Psychophysics 73:971–995.

Stecker GC, Hafter ER (2000) An effect of temporal asymmetry on loudness. The Journal of the Acoustical Society of America 107:3358–3368.

Storace DA, Higgins NC, Read HL (2011) Thalamocortical pathway specialization for sound frequency resolution. Journal of Comparative Neurology 519:177–193.

Thomas DA, Takahashi LK, Barfield RJ (1983) Analysis of ultrasonic vocalizations emitted by intruders during aggressive encounters among rats (Rattus norvegicus). Journal of comparative psychology 97:201.

Trommershauser J, Kording K, Landy MS (2011) Sensory cue integration Oxford University Press.

Wang XJ, Kennedy H (2016) Brain structure and dynamics across scales: in search of rules. Current opinion in neurobiology 37:92–98.

Wöhr M, Schwarting RK (2008) Maternal care, isolation-induced infant ultrasonic calling, and their relations to adult anxiety-related behavior in the rat. Behavioral neuroscience 122:310.

Yartsev M, Hanks T, Yoon A, Brody C (2018) Causal contribution and dynamical encoding in the striatum during evidence accumulation. Elife 7.

Yerxa T, Kee E, DeWeese M, Cooper E (2020a) Efficient sensory coding of multidimensional stimuli. PLoS Comput Biol 16:e1008146.

Yerxa TE, Kee E, DeWeese MR, Cooper EA (2020b) Efficient sensory coding of multidimensional stimuli. PLoS computational biology 16:e1008146.

Zhai X, Khatami F, Sadeghi M, He F, Read HL, Stevenson IH, Escabí Ma (2020) Distinct neural ensemble response statistics are associated with recognition and discrimination of natural sound textures. Proceedings of the National Academy of Sciences 117:31482–31493.

Zhang W, Wang H, Chen A, Gu Y, Lee T, Wong K, Wu S (2019) Complementary congruent and opposite neurons achieve concurrent multisensory integration and segregation. Elife 8.

